# Living in a multi-stressor world: nitrate pollution and thermal stress interact to affect amphibian larvae

**DOI:** 10.1101/2024.02.22.581589

**Authors:** Noa Sinai, Paula C. Eterovick, Natasha Kruger, Ben Oetken, Katharina Ruthsatz

## Abstract

The interaction of widespread stressors such as nitrate pollution and increasing temperatures associated with climate change are likely to affect aquatic ectotherms such as amphibians. The metamorphic and physiological traits of amphibian larvae during the critical onset of metamorphosis are particularly susceptible to these stressors. We conducted a common-garden experiment using *Rana temporaria* larvae subjected to four constant acclimation temperatures (18, 22, 26, 28 °C) crossed with three environmentally relevant nitrate concentrations (0, 50, 100 mg × L^-1^) to investigate the interactive and individual effects of these stressors on metamorphic (i.e., growth and development) and physiological traits (i.e., metabolism and heat tolerance) at the onset of metamorphosis. Larvae exposed to elevated nitrate concentrations and thermal stress displayed increased metabolic rates but decreased developmental rate, highlighting interactive effects of these stressors. However, nitrate pollution alone had no effect on either metamorphic or physiological traits, suggesting that detoxification processes were sufficient to maintain homeostasis but not in combination with increased acclimation temperatures. Furthermore, larvae exposed to nitrate displayed diminished abilities to exhibit temperature-induced plasticity in metamorphosis timing and heat tolerance, as well as reduced acclimation capacity in metabolic rate and heat tolerance to higher temperatures. These results highlight the importance of considering the exposure to multiple stressors when investigating how natural populations respond to global change.

## 1. Introduction

There is mounting evidence that environmental stressors associated with global change might have far-reaching consequences on organismal health and survival, which may adversely affect fitness and ultimately lead to population declines, thereby potentially contributing to ongoing global biodiversity loss (Sala et al. 2000; Díaz et al. 2019; Jaureguiberry et al. 2022). In its natural habitat, wildlife is likely to encounter multiple environmental stressors simultaneously, which often interact to yield complex ecological responses (that is, additive, antagonistic and synergistic effects; Gunderson et al. 2016; Brooks & Crowe 2019). For example, the effect of one stressor on an organism may extend to shaping the sensitivity to others, potentially compromising its ability to cope with changing environmental conditions (Rohr & Palmer 2013; Rohr et al. 2011). Therefore, to advance our understanding of the ongoing global decline in biodiversity, we need to increase our knowledge on how multiple stressors interact to affect individuals, populations, and ecosystems (Pirotta et al. 2022).

Climate change is considered to be a major driver of biodiversity loss (IPCC 2023), primarily due to its profound influence on environmental temperatures, which are one of the most important abiotic factors shaping animal distribution, ecology, and physiology (Angilletta 2009). This is of particular relevance for ectotherms, since environmental temperature determines their body temperature and thus regulates the rates of all physiological and biochemical processes that affect growth, development, and metabolism (Hochachka & Somero 1973; Angilletta et al. 2002). Consequently, climate change can impair homeostasis and other physiological mechanisms, thereby affecting organisms’ responses to co-occurring stressors (Blaustein & Kiesecker 2002; Noyes et al. 2009; Rohr et al. 2011). Moreover, climate change is often described as acting synergistically with other stressors such as environmental pollution (Rohr et al. 2011; Noyes & Lema 2015; Pulsford et al. 2019), resulting, for instance, from industrial waste, agricultural chemicals, or urban runoff (Rohr et al. 2006; Wake & Vredenburg 2008; Sigmund et al. 2023). In particular, aquatic habitats pose a sink for all kinds of contaminants including heavy metals, pharmaceuticals, nutrients, and pesticides (Rzymski et al. 2017; Mushtag et al. 2020; Kumar et al. 2021). Several studies have investigated the interactive effects of pollutants and climate change-induced thermal stress on biota, suggesting that this interaction may take several forms/directions (rev. in Rohr et al. 2011; Hooper et al. 2012): Thermal stress might increase pollutant toxicity by decreasing energy available for detoxification (i.e., temperature-induced pollutant sensitivity; Osterauer & Kohler 2008; Noyes et al. 2009). Alternatively, pollutants might decrease tolerance to thermal stress by reducing energy available for temperature regulation or acclimation (i.e., pollutant-induced temperature sensitivity; Lannig et al. 2006; Noyes et al. 2009) or by acting as endocrine disruptors affecting hormonal regulations necessary for acclimation such as thyroid hormones (TH) (Little & Seebacher 2014; Little 2021). However, interactive effects of multiple stressors remain difficult to predict and existing studies often oversimplify possible interactions between stressors (rev. in Ruthsatz & Glos 2024) or do not address the complexity of factors or their multitude of interactive effects, focusing instead on direct effects of single factors (Walls & Gabor 2019).

Given their dramatic population declines, the need to study interactions among multiple stressors seems particularly compelling for amphibians (Sih et al. 2004; Walls & Gabor 2019). They are directly or indirectly affected by various stressors throughout their lives, suggesting that the causes for their population declines are complex and include interactions of multiple stressors (Blaustein & Kiesecker 2002; Hayes et al. 2010; Blaustein et al. 2011). Recently, climate change and environmental pollution have been identified as major drivers of global amphibian population declines (Luedtke et al. 2023), emphasizing the need for understanding how these stressors interact to impair amphibian health and survival. Amphibians may be particularly sensitive to interactions between climate change and pollutants due to their highly permeable skin, which offers little resistance to evaporative water loss or uptake of pollutants, and as ectotherms, amphibians may be sensitive to temperature changes (Navas et al. 2008; rev. in Larsen 2021). In fact, several studies indicate that exposure to pollutants and the effects of climate change may be worse for amphibians than either stressor alone (e.g., Boone & Bridges 1999, Rohr & Palmer 2005; Baier et al. 2016; Ruthsatz et al. 2018a, 2020a, 2023-MP; but not: Rohr et al. 2011). Despite possible additive or synergistic effects, interactions between pollution and climate change may also affect amphibians’ ability to cope with changing environments through phenotypic plasticity (Ruthsatz et al. 2020a). This is especially important for amphibian larvae as they may experience greater exposure to multiple stressors in the aquatic habitat compared to terrestrial life stages (Rohr & Palmer 2005). Moreover, they often rely on developmental plasticity to cope with changing environmental conditions, as these stages cannot easily escape an unfavorable habitat (Newman 1992; Merilä et al. 2000; Gomez-Mestre et al. 2013). However, studies assessing whether and how environmental stressors might affect the coping capacity of (larval) amphibians through phenotypic plasticity are still rare.

Amphibian breeding habitats are often found in agricultural areas and the breeding season of many species coincides with arable farming activities (Brühl et al. 2013; Goessens et al. 2022). Therefore, exposure to agrochemicals, such as pesticides and fertilizers, can have a significant ecological impact on amphibians across life stages (rev. in Mann et al. 2009; Baker et al. 2013). Agrochemicals are receiving increasing attention as a potential cause of amphibian declines, either acting alone or in combination with other stressors such as those associated with climate change (rev. in Ruthsatz & Glos 2024). While the potential organismal and population-level effects of a variety of pesticides on amphibians as non-target organisms have been extensively studied (Van Meter et al. 2019; Trudeau et al. 2020), relatively little is known about the effects of fertilizers on amphibian health, especially in combination with climate change (rev. in Ruthsatz et al. 2022a). As the major components of fertilizers, nitrate (NO_3_**^-^**) is an ubiquitous pollutant in habitats of (larval) amphibians (e.g., Rouse et al. 1999; De Wijer et al. 2003; Ortiz-Santaliestra & Sparling 2007) and has been shown to affect growth (Garriga et al. 2017), development (Wang et al. 2015; Xie et al. 2019), reduce survival (Ruthsatz et al. 2022a), and has also been linked to disruption of the thyroid hormone (TH) system during metamorphosis (Wang et al 2015; Poulsen et al. 2018; but not: Edwards et al. 2006). Given that THs play a key regulatory role in the onset and timing of metamorphosis in amphibians (rev. in Shi 2000; Denver 2021) and thermal acclimation in ectotherms in general (Little 2021), nitrate exposure may not only alter growth and development, but also reduce the ability to cope with new thermal challenges by exhibiting developmental or physiological plasticity. However, it remains unclear whether nitrate exposure impairs amphibian responses to climate change.

Here, we studied the interactive effects of thermal stress and nitrate pollution on larvae of the European common frog (*Rana temporaria*) in a laboratory experiment. To that end, we raised larvae at four, constant acclimation temperatures (18, 22, 26, 28 °C) crossed with three environmentally relevant nitrate concentrations (0, 50, 100 mg × L^-1^). Specifically, we examined whether and how nitrate exposure and acclimation temperature interact to affect metamorphic (i.e., growth and development) and physiological traits (i.e., metabolism and heat tolerance) of amphibian larvae at the onset of metamorphosis. Furthermore, we investigated whether exposure to nitrate pollution impairs the capacity for temperature-induced plasticity in developmental rate as well as the capacity for physiological acclimation in standard metabolic rate (SMR) and heat tolerance (i.e., CT_max_). We predicted that thermal stress might exacerbate the effects of nitrate pollution on metamorphic and physiological traits by decreasing energy available for detoxification (Osterauer & Kohler 2008; Noyes et al. 2009), resulting in ramifications for growth, development, metabolism, and heat tolerance (Ruthsatz et al. 2020a,b,c). Also, we predicted that exposure to nitrate pollution might decrease tolerance to thermal stress since nitrate might disrupt hormone signaling necessary for development/timing of metamorphosis and acclimation (Little & Seebacher 2014; Little 2021). Furthermore, detoxification processes might reduce energy available for plastic responses in developmental rate and thermal acclimation (Lannig et al. 2006; Noyes et al. 2009). Our study aimed to enhance our understanding of how multiple-stressor environments may increase the vulnerability of amphibians to global change and population declines.

## 2. Materials and Methods

### 2.1 Study species field sampling

*Rana temporaria* was chosen as a model species as its capacity for phenotypic plasticity in metamorphic (Lind & Johansson 2007; Merilä et al. 2004) and physiological traits (Ruthsatz et al. 2022a,b) in response to variation in environmental factors is known to be relatively high compared to other amphibian species (Laurila & Kujasalo 1999; Vences et al. 2017; Sinai et al. 2022). Five clutches of the European common frog (*Rana temporaria*) were collected at the locality Kleiwiesen (52.328 N, 10.582 E), a site near Braunschweig, Lower Saxony.

### 2.2 Experimental setup and animal husbandry

The experiment was conducted in a climate chamber (Kälte-Klimatechnik-Frauenstein GmbH, Germany) with a 14:10 h light:dark cycle at the Zoological Institute at Technische Universität Braunschweig. Egg clutches were kept separately until hatching in five 12-L plastic buckets filled with 5 L of fresh pond water at 14 ± 0.2 °C. Freshwater was obtained from the original pond every three days and stored at 4 °C. Approximately one-third of the water volume in the buckets was replaced with the stored pond water (after reaching 14°C) each day to prevent excessive waste accumulation. After hatching, air temperature in the climate chamber was increased to 18 ± 0.1 °C within 24 h and larvae were allowed to develop to stage 25 (free-swimming larvae; Gosner 1960) before allocation to experimental treatments. Three larvae from each clutch were then randomly selected and allocated to 15 standard 12-L glass aquaria filled with 9 L of aged de-chlorinated tap water at four different temperature regimes (18 ± 0.2 °, 22 ± 0.3 °, 26 ± 0.4 °, 28 ± 0.3 °C) crossed with three levels of nitrate exposure (0, 50, 100 mg × L^-1^) in a common-garden experiment using three replicates for each temperature x nitrate treatment combination (4 temperatures **×** 3 nitrate treatments **×** 3 replicates = 36 aquaria). Each aquarium housed 15 larvae (15 larvae **×** 36 aquaria = 540 larvae in total), which resulted in a larval density of 1.66 larvae × L^-1^ in the beginning of the experiment. Water temperatures were achieved by adjustable heating elements (JBL PROTEMP S 25, Germany). Continuous water aeration was provided through air stones connected to aquaria pumps. Water was changed every third day keeping the respective rearing temperature to avoid any temperature fluctuations.

When larvae reached Gosner stage 38 (i.e., all toes separated; Gosner 1960), three larvae per aquarium were randomly picked, removed from the aquaria, and euthanized for another study (Eterovick et al. 2024). The remaining larvae were kept in the aquaria for the present study. When larvae were approaching the onset of metamorphosis, tanks were surveyed daily to retrieve metamorphosing individuals (i.e., at forelimb emergence, Gosner stage 42; Gosner 1960). Metamorphs (Gosner stage 42) were removed from aquaria and assigned to either oxygen consumption or thermal tolerance assessments. The experiments ran for 5 weeks, all surviving larvae needed from 13 to 34 days to reach the onset of metamorphosis, the terminal sampling point in the present study. After completion of physiological assessments, metamorphs were kept individually in 470 mL lidded plastic containers with 2 mm of aged de-chlorinated tap water at the bottom at an air temperature of 18 ± 0.1 °C until they completed tail resorption (Gosner stage 46, Gosner 1960) and were used in other studies.

Larvae were fed 50 % high-protein powdered fish food (Sera micron breeding feed for fish and amphibians, Sera, 52518 Heinsberg, Germany) and 50% spirulina algae. *Ad libitum* rations were provided twice a day to guarantee that food was available in abundance. The size of the rations was continuously adjusted to account for changes in the size of larvae and the number of individuals in each aquarium.

### 2.3 Nitrate exposure

Nitrate concentrations were achieved using sodium nitrate stock solutions (NaNO_3_; e.g., Oromí et al. 2009; Wang et al. 2015) in accordance with Ruthsatz et al. (2022a). Briefly, stock solutions of reagent-grade sodium nitrate (>99% pure; Carl Roth, 76185 Karlsruhe, Germany) were prepared with autoclaved water and the volumes required to achieve target concentrations were pipetted into the aquaria (i.e., 0, 50, 100 mg × L^-1^). After each water change, the procedure was repeated to achieve the target nitrate concentration again. The nitrate concentrations selected for the experiment are within the environmental ranges observed in both surface and ground waters in Germany (Sundermann et al. 2020) and align with the environmental ranges measured in water bodies where amphibians breed (De Wijer et al. 2003; Rouse et al. 1999; Johansson et al. 2001).

### 2.4 Natal pond and experimental water parameters

Water quality was monitored before each water change during the experiment with the use of the AQUA-Check 2 photometer (Söll GmbH, 95030 Hof, Germany; N = 12). Natal pond water parameters were determined at the time of clutch collection and every time we collected fresh pond water. Three water samples each were taken from the edge and the middle of the pond and pooled for each analysis. Measurements included nitrate (NO_3_^-^ ), nitrite (NO_2_^-^ ), ammonium (NH_4_^+^), pH, phosphate (PO_4_^3-^)-, copper (Cu^2+^), iron (Fe), and lead (Pb) in mg **×** L^-1^ (Table S1).

### 2.5 Life history variables

To determine developmental stage, individuals were visually inspected one day prior to sampling with bare eye or examined under a binocular microscope (KeyenceX-500). Ontogenetic stage was determined by evaluating the status of key morphological features typical of specific stages, as detailed in Gosner (1960). The age was given by the number of days after hatching (dah).

We measured total length (TL), snout-vent-length (SVL), and body mass at Gosner stage 25 (i.e., at the start of the experiment, 4 dah) and at the onset of metamorphosis (i.e., at forelimb emergence, Gosner stage 42; Gosner 1960) after finishing the physiological measurements (Section 2.6). TL and SVL were measured with a caliper to the nearest 0.5 mm. To measure body mass, specimens were dry blotted and weighed to the nearest 0.001 g with an electronic balance (Sartorius A200 S, Germany). Growth rate (mg × d^-1^) was given by the individual difference in mass between Gosner stage 25 and metamorphosis divided by the number of days needed to reach metamorphosis from Gosner stage 25.

### 2.6 Physiological measurements

As animals reached the onset of metamorphosis, we allocated them to either respiration measurements or heat tolerance assessments. After taking physiological measurements, we determined the SVL and blotted wet body mass of each animal. Then, animals were placed back into their rearing plastic containers.

#### 2.6.1 Metabolism measurements

Oxygen consumption was measured in six randomly chosen larvae from each aquarium (N_total_=216) using closed respirometry during the natural activity phase between 0830 and 2130 h (Orlofske & Hopkins 2009). To that end, larvae were placed in 30-mL glass vessels filled with autoclaved tap water to exclude microbial oxygen consumption and sealed with airtight rubber plugs. Water contained treatment-level concentrations of nitrate as experienced by individuals during development. In each vessel, a chemical optical oxygen sensor spot was integrated, which was connected to a multichannel oxygen measuring system (Oxy-4 SMA; PreSens Precision Sensing GmbH, Regensburg, Germany) via a fiber optic sensor (Polymer Optical Fiber POF, PreSens Precision Sensing GmbH, Regensburg, Germany). A temperature probe submerged in water at the same temperature as the respiration vials provided temperature compensation for dissolved oxygen measurements. No fasting prior to the respiratory measurements was needed because larvae stop feeding due to the remodeling of mouthparts and digestive tract during metamorphosis (Hourdry et al. 1996; Ruthsatz et al. 2019,2020b).

Measurements started 10 min after allocating the animals to the vessels and O_2_ concentration was recorded every 15 s and measured as ml O_2_ **×** L^-1^ (volume not corrected for the size of the animal). Prior to each trial, the O_2_ fiber optic sensors were calibrated using air-saturated water and a factory-set zero oxygen calibration point at the respective measuring. Water temperature was controlled by the continuous mixing of the water bath. Oxygen consumption was measured for every animal for 30 min. Empty (control) chambers were run simultaneously in every trial and values were adjusted accordingly. We ensured that less than 10 % of total O_2_ was removed during any measurement period to avoid changes in respiration at low O_2_ saturation levels.

To determine acclimation capacity in oxygen consumption in response to temperature, we measured oxygen consumption not only at acclimation temperature but also at acclimation temperature + 2 °C and + 4 °C. Thereto, the water bath temperature was increased at a rate of + 0.1 °C × min^-1^ following the initial measurement. During this ramping period, the glass vessels were open in order to allow aeration. The animals were given a 10-min period to acclimate to the target temperature before measurements were taken for 30 min. This procedure was repeated for the second temperature step. After taking the final measurements, the temperature of the water bath was gradually decreased at a rate of - 0.1 °C × min^-1^ until the larvae reached their acclimation temperature. All larvae survived the metabolism measurements (i.e., no mortality after 24 h).

##### 2.6.1.1 Standard metabolic rate calculations

Prior to statistical analysis, we plotted O_2_ consumption of each animal over time and visually assessed activity peaks to exclude them for the determination of standard metabolic rate (SMR). Readings generated in the first 5 min of each assay were discarded to remove acute effects of handling. The SMR was expressed in ml O_2_ **×** h^-1^ **×** g^-1^ dry-blotted body mass and was determined from the slope of linear least squares regression of O_2_ concentration vs. time (Hastings & Burggren, 1995; Ruthsatz et al. 2018a, 2019). The analyses were performed in PreSens Oxygen Calculator Software (PreSens Precision Sensing GmbH, Regensburg, Germany) as well as Microsoft Excel following our procedure explained in Ruthsatz et al. (2022a).

#### 2.6.2 Heat tolerance

Critical thermal maximum (CT_max_) was measured as a proxy for heat tolerance and evaluated in a minimum of three and a maximum of six larvae per aquarium (i.e., remaining larvae not assigned to the respiration measurements; N_total_=169) using the dynamic method according to Cowles and Bogert (1944) and Hutchison (1961), except for the endpoint (Wu & Kam 2005). The critical thermal maximum was defined as the temperature at which locomotor activity became disorganized and the animals lost their righting response (Holzman & McManus 1973; Lutterschmidt & Hutchison 1997). The righting response is ecologically relevant for selection on thermal traits as an animal that is unable to use its righting reflex is unable to escape predators (von May et al. 2019). To assess this endpoint, the animal is flipped on the back in the water with a probe (Wu & Kam 2005; Turriago et al. 2015; Ruthsatz et al. 2020c).

Each individual was placed in a 250-mL beaker filled with 200 ml of aerated water, which was then placed in a temperature-controlled water bath. Water contained treatment-level concentrations of nitrate as experienced by individuals during development. The initial temperature in the water bath was set at the individual’s acclimation temperature. Water bath temperature was increased starting at the respective acclimation temperature at a rate of + 0.1°C × min^-1^ (Lutterschmidt & Hutchison 1997). Heat tolerance measurements were carried out between 1200 and 1700 h to control for the influence of natural circadian rhythms and photoperiod (Agudelo-Cantero & Navas 2019). Furthermore, animals experience the highest diurnal temperatures in their natural habitats during this period.

When animals reached their CT_max_, we removed them gently from the beakers and transferred them to a beaker filled with water at

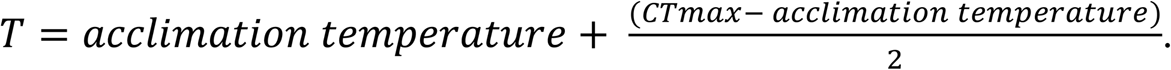

The beaker was placed in a water bath set to the appropriate acclimation temperature to allow for a gradual decrease in body temperature until the water in the beaker reached the same temperature as the water bath. All larvae recovered from the CT_max_ assessments (i.e., no mortality after 24 h).

### 2.7 Statistics

For all statistical tests Cran R (Version 4.3.2, R Development Core Team 2021) for Windows was used and significance was set at α<0.05.

#### 2.7.1 Plasticity in developmental rate

To determine the thermal reaction norm (i.e., sensitivity) of developmental rate to temperature variation, we performed single linear regressions of acclimation temperature (independent variable) and age at the onset of metamorphosis (dependent variable) within each nitrate treatment for each aquarium independently. The slope of each regression describes the change in age at metamorphosis with a given change in developmental temperature and was used as a plasticity index (PIX) according to Ruthsatz et al. (2018b) and Sinai et al. (2022). Higher absolute values of PIX correspond to higher plasticity (e.g. greater sensitivity of developmental rate to temperature).

#### 2.7.2 Acclimation capacity in SMR

Acclimation capacity in SMR to short-term temperature variation (i.e., acute thermal sensitivity) can be identified by calculating the coefficient Q_10_, which is a measure of the rate of change of the metabolic processes by an increase of temperature of 10 °C (e.g., Dalvi et al. 2009). The relative change in respiration rate over a 10 °C interval (Q_10_) was calculated using the formula:

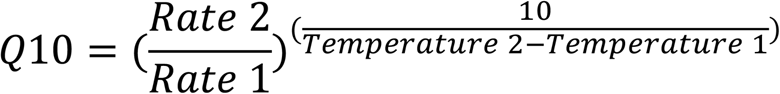

We calculate the mean Q_10_ values from the Q_10_ values between the differences within each nitrate treatment. The highest Q_10_ values would represent the greatest increases in SMR (in this case, oxygen consumption) at higher temperatures and *vice versa*.

#### 2.7.3 Thermal tolerance plasticity

A well-established method to measure acclimation capacity in thermal tolerance in ectothermic animals is the calculation of the acclimation response ration (ARR), i.e., the slope of the linear function describing the change in thermal tolerance with a given change in acclimation temperature (Hutchison 1961; Claussen 1977). We calculated the ARR for CT_max_ within each aquarium using the equation according to Claussen (1977):

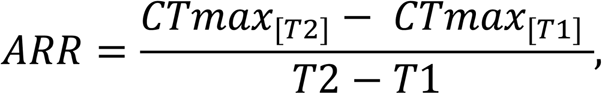

where T represents the acclimation temperature (°C; with T2 = highest acclimation temperature, 28°C, and T1 = lowest acclimation temperature, 18 °C) and CT_max_ the heat tolerance estimates (°C). Higher absolute values of ARR correspond to higher plasticity in thermal tolerance limits (i.e., greater acclimation capacity; Claussen 1977; Kingsolver & Huey 1998). An acclimation response ratio of 1.00 indicates a 100 % acclimation in thermal tolerance to a temperature increase of 1 °C (Morley et al. 2019).

#### 2.7.4 Effects of temperature and nitrate pollution on metamorphic and physiological traits

We tested the effect of “acclimation temperature” (i.e., respective rearing temperature) and “nitrate treatment” as fixed factors on metamorphic traits (age, SVL, growth rate), SMR, heat tolerance (measured as CT_max_), and survival in six separate linear mixed models (LMMs) using the *lmer()* function in the lme4 package (https://cran.r-project.org/package=lme4) with REML restricted maximum likelihood) as method for parameter estimation (Bates et al. 2010). “Body mass” (Leiva et al. 2019; Ruthsatz et al. 2022b) was included as covariate to LMM on heat tolerance. To investigate whether acclimation temperature and nitrate pollution interact to affect growth, development, and physiology, we also included the interactive effect of “acclimation temperature” and “nitrate treatment” as fixed factor. Residuals of each model were visually checked for normal distribution (QQ-plots). To address dependencies in the data and to avoid pseudoreplication, “aquarium” was included as a random factor in all models. N refers to the total number of animals and n to the total number of aquaria.

#### 2.7.5 Effects of nitrate pollution on thermal acclimation capacity

To investigate whether nitrate pollution affects the capacity for thermal acclimation in amphibian larvae, data were analyzed using three separate LMMs using the *lmer()* function (Bates et al. 2010) in the lme4 package (https://cran.r-project.org/package=lme4) with REML restricted maximum likelihood) as method for parameter estimation (Bates et al. 2018). As dependent variables, we used the PIX of developmental rate, mean Q_10_ of SMR, and the ARR of CT_max_ in three separate models. “Nitrate” treatment was included as a fixed factor in all models. “Body mass” was included as covariate in the model on ARR since CT_max_ analyses revealed a significant effect of body mass (present study; Ruthsatz et al. 2022b). To avoid pseudoreplication, the categorical predictor “aquarium” was included as a random factor in all models. LMMs were followed by post hoc comparisons with Bonferroni correction (Tukey’s test; Tukey’s HSD function, multcomp package) to compare all possible pairwise combinations of nitrate treatment when overall tests were significant.

## 3. Results

### 3.1 Effects of temperature and nitrate pollution on metamorphic and physiological traits

#### Survival

Survival decreased with temperature and nitrate concentration (Table S2; Table 1). Furthermore, within each temperature treatment, survival decreased as nitrate levels increased. Thus, survival was highest at 18 °C and at 0 mg **×** L^-1^ nitrate (100 % ± 0) and lowest at 28 °C and at 100 mg **×** L^-1^ (83.33 % ± 0).

**Table 1.**
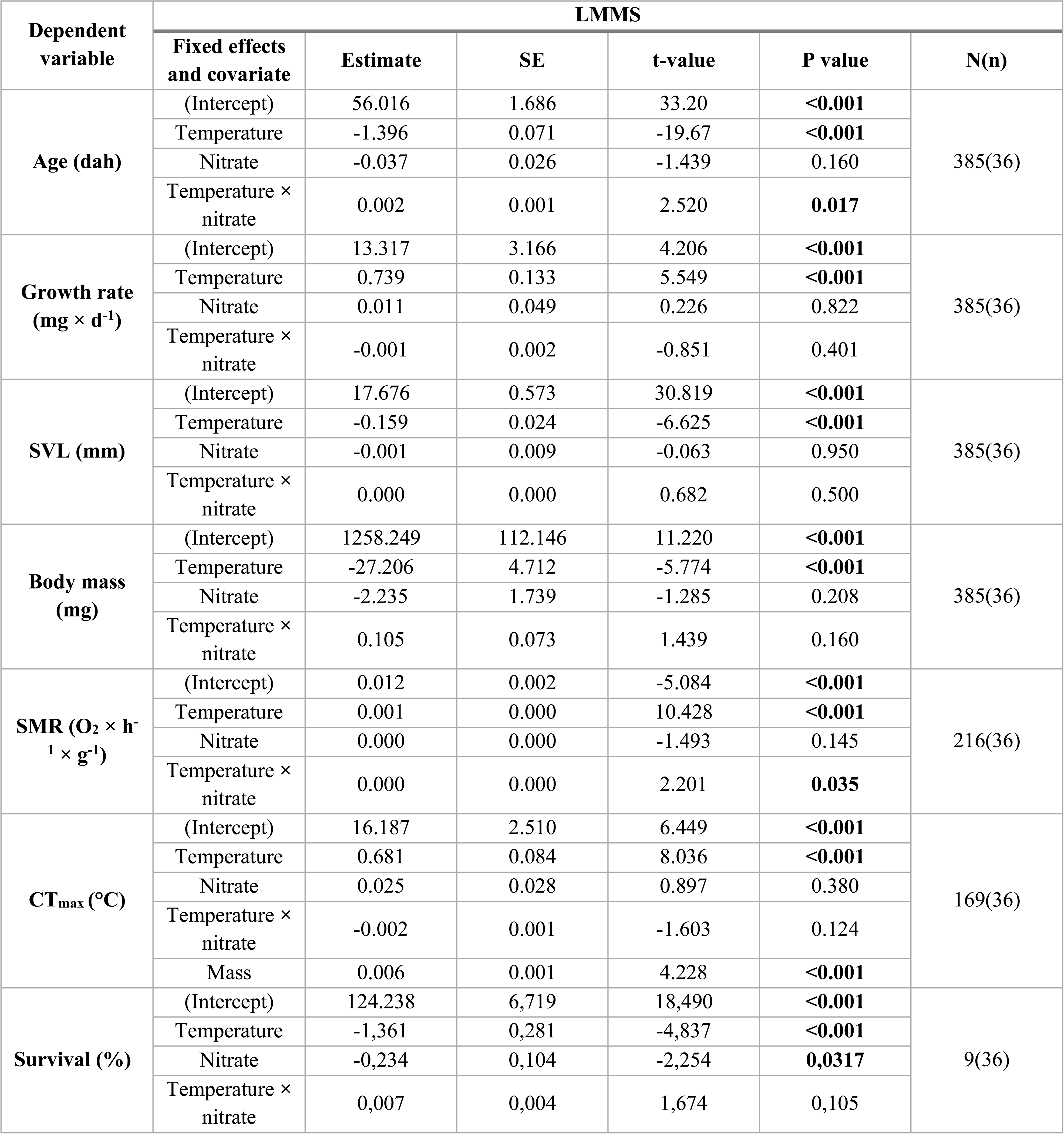
Effects of acclimation temperature, exposure to nitrate pollution, and their interaction on metamorphic and physiological traits of the European common frog (*Rana temporaria*) at the onset of metamorphosis (Gosner stage 42; Gosner 1960). Results were obtained from separate LMMs, using “temperature” and “nitrate” treatments as well as the interactive effect of both as fixed factors; “aquarium” as random factor; “mass” as the covariate in the model on CT_max_. N is the total number of analyzed data points (i.e., individual larvae) across treatments; n refers to the total number of analyzed aquaria. Significance was set at p<0.05. Abbreviations: LMMs, linear mixed models. SMR = standard metabolic rate (O_2_ **×** h^-1^ **×** g^-1^). CT_max_ = critical thermal maximum/heat tolerance (°C). Age in days after hatching (dah); growth rate = increase in body mass during the course of the experiment (mg **×** d^-1^).

#### Metamorphic traits

Acclimation temperature significantly affected age and size (mass and SVL) at metamorphosis as well as growth rate during larval development (Figure 1; Table 1). Whereas age and size decreased with temperature, a higher acclimation temperature resulted in a higher growth rate. In contrast, nitrate exposure had no effect on metamorphic traits. Acclimation temperature and nitrate exposure interacted to affect age at metamorphosis, with animals at higher temperatures (26 and 28 °C) exposed to 100 mg **×** L^-1^ revealing a significantly higher age than at lower nitrate concentrations.

**Figure 1.**
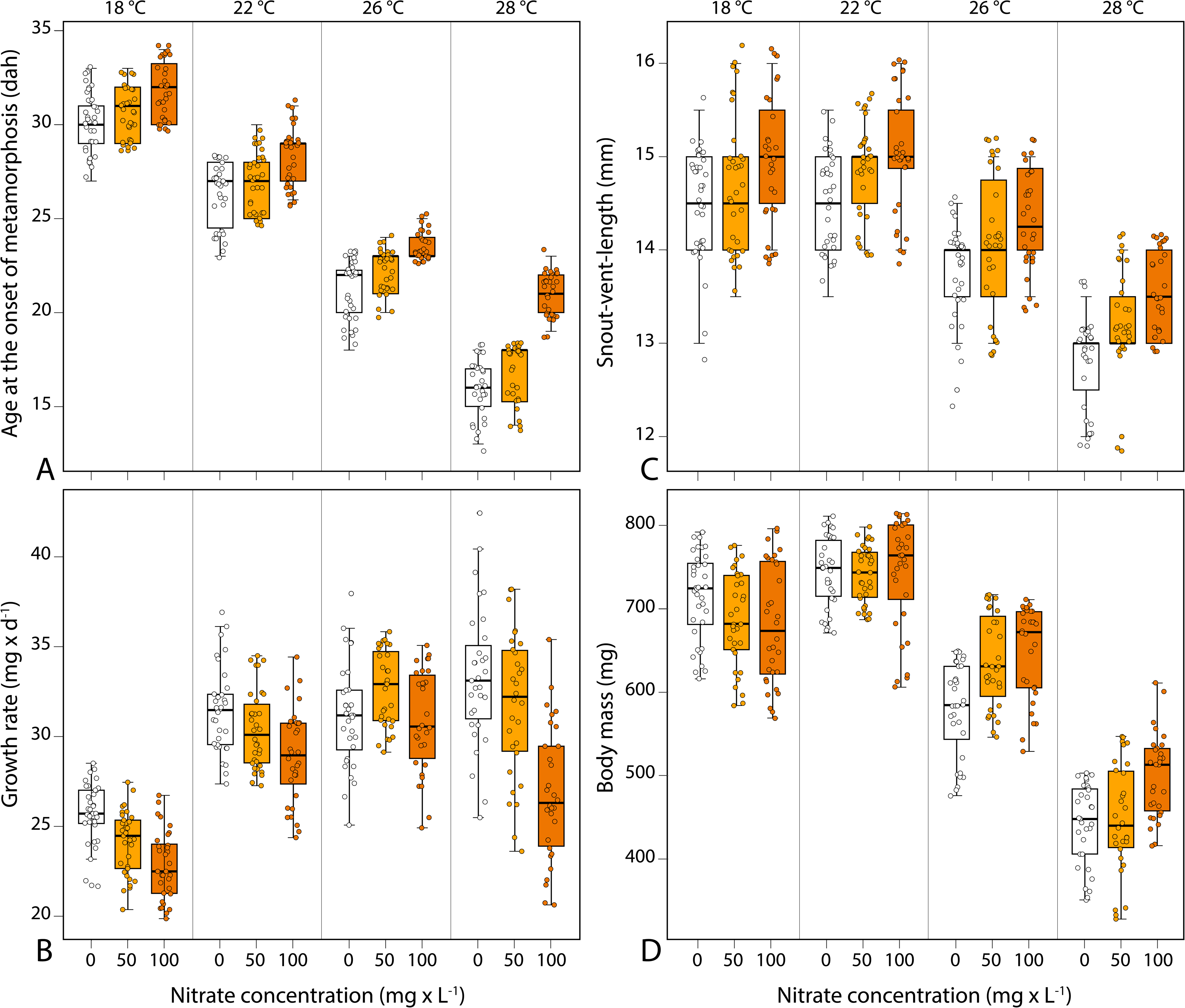
Effects of acclimation temperature, exposure to nitrate pollution, and their interaction on metamorphic traits of the European common frog (*Rana temporaria*) at the onset of metamorphosis (Gosner stage 42; Gosner 1960). **A** Age (days after hatching, dah). **B** Growth rate (mg × d^-1^). **C** Snout-vent-length (mm, SVL). **D** Body mass (mg).

#### Physiological traits

Standard metabolic rate and heat tolerance significantly increased with acclimation temperature, whereas nitrate exposure had no effect on both physiological traits (Table 1; Figure 2). Acclimation temperature and nitrate exposure had a significant interactive effect on SMR but not on heat tolerance. SMR increased stronger with increasing nitrate at higher temperatures.

**Figure 2.**
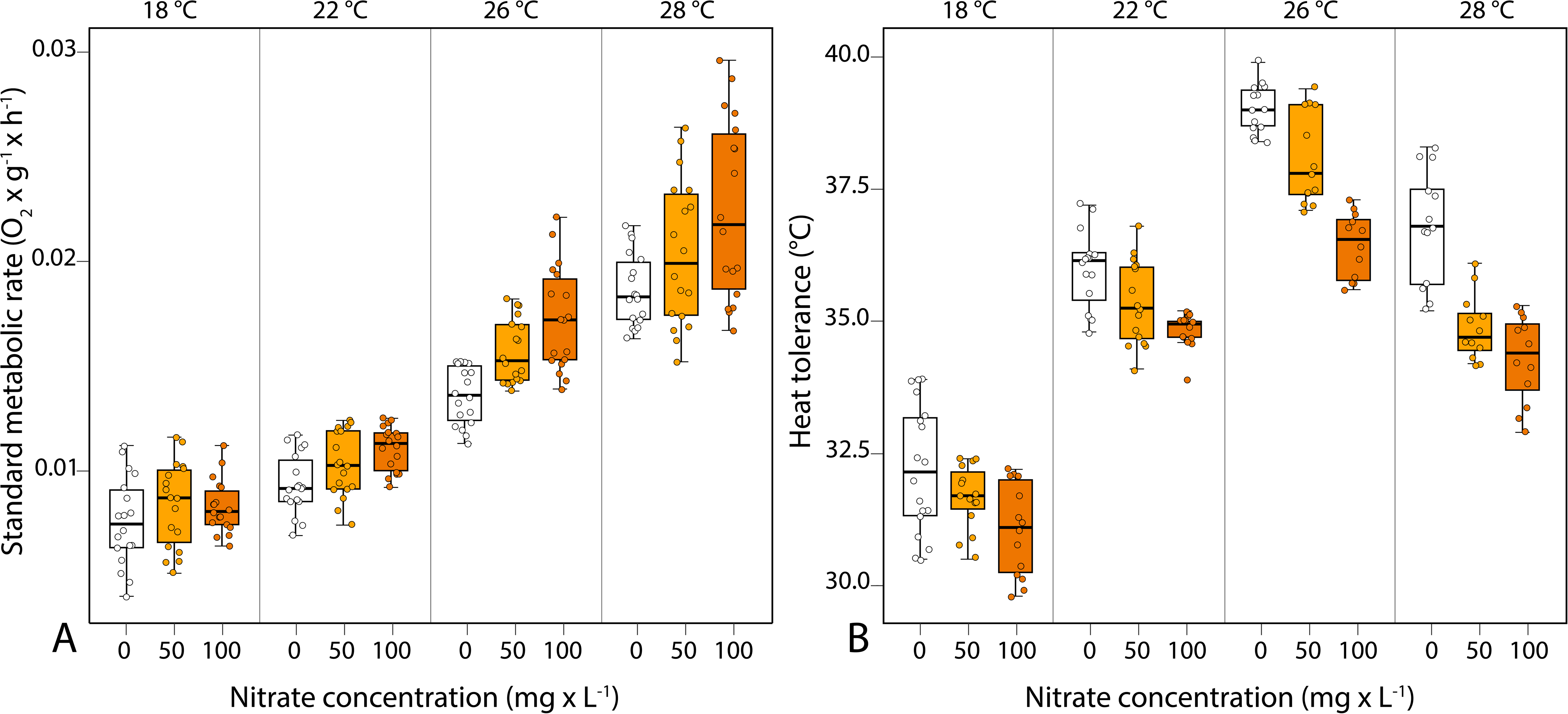
Effects of acclimation temperature, exposure to nitrate pollution, and their interaction on physiological traits of the European common frog (*Rana temporaria*) at the onset of metamorphosis (Gosner stage 42; Gosner 1960). **A** Standard metabolic rate (O_2_ × h^-1^ × g^-1^, SMR). **B** Heat tolerance measured as critical thermal maximum (°C, CT_max_).

### 3.2 Effects of nitrate pollution on thermal acclimation capacity

Nitrate exposure significantly reduced the capacity for thermal acclimation in SMR, heat tolerance, and temperature-induced developmental plasticity as indicated by a higher Q_10_, and lower ARR and PIX values (Table 2; Figure 3). Pairwise comparisons revealed significant differences in the capacity for temperature-induced plasticity in developmental rate between the control group and the nitrate concentration of 100 mg × L^-1^ as well as between 50 and 100 L^-1^. Q_10_ was significantly different between all nitrate treatments. In contrast, ARR was significantly lower when animals were exposed to nitrate but independent of the concentration.

**Figure 3.**
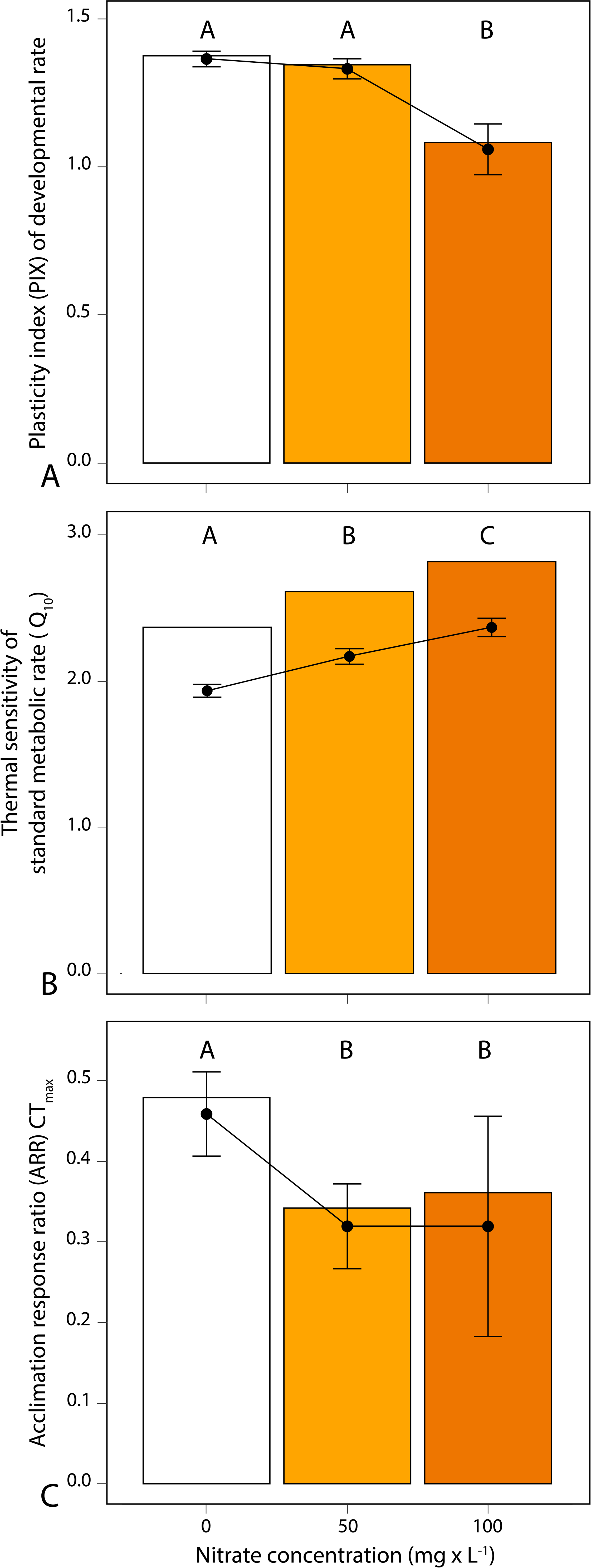
Effects of nitrate pollution on the thermal acclimation capacity in the European common frog (*Rana temporaria*) at the onset of metamorphosis (Gosner stage 42; Gosner 1960). **A** Plasticity index (PIX) for temperature-induced plasticity in developmental rate (N=36). **B** Mean thermal sensitivity of standard metabolic rate (Q_10_) (N=36). **C** Acclimation response ratio (ARR) of heat tolerance measured as critical thermal maximum (CT_max_). Error bars represent means ± 95% confidence intervals (N=216). Different letters above the columns indicate significant differences between the treatments.

**Table 2.** Effects of nitrate pollution on the capacity for thermal acclimation in developmental rate, standard metabolic rate, and heat tolerance in the European common frog (*Rana temporaria*) at the onset of metamorphosis (Gosner stage 42; Gosner 1960). Results were obtained from separate LMMs, using “nitrate” treatment as the fixed factor; “aquarium” as a random factor. Pairwise multiple comparisons between nitrate treatments (0, 50, and 100 mg × L^-1^) using Tukey’s Test as a post hoc test with Bonferroni correction. N = total number of analyzed data points, n = total number of aquaria analyzed. Significance was set at p < 0.05. Abbreviations: LMMs, linear mixed models. PIX = plasticity index of developmental rate. Mean Q_10_ = thermal sensitivity of standard metabolic rate. ARR = acclimation response ratio; CT_max_ = critical thermal maximum/heat tolerance. See text for further details on the plasticity indices.

| Dependent variable<br>N(n) | LMMs |  |  |  |  | Post hoc test for pairwise multiple comparisons |  |  |  |  |
| --- | --- | --- | --- | --- | --- | --- | --- | --- | --- | --- |
|  | Fixed effects | Estimate | SE | t-value | P value | Pairwise comparison | Estimate | SE | z-value | P value |
| PIX<br>N(n)=9(36) | (Intercept) | 1.402 | 0.035 | 39.764 | <0.001 | 0-50 | -0.033 | 0.017 | -1.859 | 0.151 |
|  | Nitrate | -0.003 | 0.001 | -5.578 | <0.001 | 0-100 | -0.304 | 0.017 | -17.024 | <0.001 |
|  |  |  |  |  |  | 50-100 | -0.271 | 0.017 | -15.165 | <0.001 |
| Mean $Q_{10}$<br>N(n)=216(36) | (Intercept) | 1.945 | 0.041 | 48.027 | <0.001 | 0-50 | 0.235 | 0.064 | 3.702 | <0.001 |
|  | Nitrate | 0.004 | 0.001 | 6.927 | <0.001 | 0-100 | 0.435 | 0.064 | 6.835 | <0.001 |
|  |  |  |  |  |  | 50-100 | 0.199 | 0.0634 | 3.133 | 0.014 |
| ARR $\text{CT}_{\text{max}}$<br>N(n)=9(36) | (Intercept) | 0.435 | 0.026 | 16.506 | <0.001 | 0-50 | -0.140 | 0.029 | -4.723 | <0.001 |
|  | Nitrate | -0.001 | 0.000 | -3.409 | 0.011 | 0-100 | -0.140 | 0.029 | -4.723 | <0.001 |
|  |  |  |  |  |  | 50-100 | -0.000 | 0.029 | 0.000 | 1 |

## 4. Discussion

Given increasing concerns over ongoing global climate change, there is an urgent need to develop a better understanding of the interactive effects of multiple stressors on wildlife. Several studies have shown that the combination of pollution stress with other environmental stressors, such as heat stress or desiccation, can often result in a stronger-than-additive effect on organisms (e.g., Holmstrup et al. 2010; Rohr et al. 2011; Moe et al. 2013; Noyes & Lema 2015; Wu & Seebacher 2021). Here, we find an interactive effect of ecologically relevant nitrate concentrations and thermal stress on both development and metabolism, alongside impacts of nitrate pollution on thermal acclimation capacity. Larvae subjected to elevated acclimation temperatures and nitrate levels revealed increased metabolic rates but a decrease in developmental rate. Furthermore, larvae exposed to nitrate displayed diminished abilities to exhibit temperature-induced plasticity in metamorphosis timing and heat tolerance, as well as a reduced acclimation capacity in metabolic rate and heat tolerance to higher temperatures. The present study therefore underscores that multiple-stressor environments may increase the vulnerability of amphibians to global (climate) change.

We hypothesized that higher acclimation temperatures and nitrate exposure would interact to affect metamorphic and physiological traits. Larvae exposed to higher acclimation temperatures and nitrate levels would show a decrease in growth, development, and heat tolerance but an increase in metabolic rate. We did not observe an interactive effect of acclimation temperature and nitrate on growth and heat tolerance, but it caused a decrease in developmental rate and an increase in metabolic rate. Notably, nitrate pollution alone had no effect on either metamorphic or physiological traits, suggesting that detoxification processes were sufficient to maintain homeostasis but not at increased acclimation temperatures. In a recent study on larvae of *R. temporaria* we observed that nitrate exposure induced changes in gut microbiome composition that were associated to a predicted increase in a bacteria-associated nitrate reduction metabolic pathway (Eterovick et al. 2024), what likely facilitates denitrification/detoxification processes (Tiso & Schechter 2015; Lo et al. 2022). In combination with thermal stress (i.e., higher acclimation temperatures), however, the capacity of the gut microbiome to buffer detrimental effects of nitrate pollution on larval health was hampered (Eterovick et al. 2024). The impairment of this adaptive coping capacity with nitrate pollution could be behind the interactive effects of acclimation temperature and nitrate exposure also observed here. Moreover, the increased metabolic rate found in larvae exposed to both stressors is likely to increase energy demands for both maintenance and development (Orlofske & Hopkins 2009; Ruthsatz et al. 2019), potentially reducing energy available for detoxification (Osterauer & Kohler 2008; Noyes et al. 2009). Further research is needed to understand the mechanisms behind the interactive effect of changing thermal conditions related to climate change on the toxicity of pollutants. Ideally, this research should address the characteristics of different pollutants to avoid oversimplification of the interaction between pollution and climate change.

Although there is a growing body of research demonstrating that changes in thermal conditions associated with climate change may amplify the adverse effects of amphibian exposure to pollutants, other studies argue that projected increases in temperature may accelerate the rate of development of amphibian larvae, especially if accompanied by pond drying (Burraco & Gomez-Mestre 2016; Ruthsatz et al. 2018b; Albecker et al. 2023), resulting in a shorter larval period and thereby reducing exposure to aquatic pollutants during the larval stage. However, our results indicate that some pollutants might reduce the capacity for temperature-induced plasticity in developmental rate as we found this capacity to be reduced in larvae exposed to nitrate. Furthermore, exposure to nitrate pollution reduced the capacity for acclimation in metabolic rate and heat tolerance to changing thermal conditions. Therefore, amphibian larvae exposed to nitrate pollution might be more vulnerable to the challenges posed by changing environmental temperatures because (1) their ability to emerge earlier from the larval habitat by accelerating their development is impaired, (2) their energy expenditure is higher due to the reduced capacity to avoid a strong increase in metabolic rate with increasing temperature, and (3) their tolerance to extreme temperatures is reduced. Mechanistically, our findings may be explained by disrupted thyroid signaling. Thyroid hormones (TH) are important for regulating metabolism, growth, and development (Shi 2000; Mullur et al. 2014; Kirschman et al. 2017) and play a crucial role in thermal acclimation (Little & Seebacher 2014; Little 2021). Disruption of thyroid function and thus, TH level by nitrate exposure can therefore interfere with physiological performance and timing of metamorphosis. In previous studies, we found that a manipulation of the TH level indeed resulted in the same effect on the capacity for temperature-induced plasticity in developmental rate (Ruthsatz et al. 2020a), metabolic rate (Ruthsatz et al. 2020b,c), and thermal tolerance (Ruthsatz et al. 2020c), supporting our hypothesis particularly considering that nitrate has also been associated with disruption of the TH system (Wang et al. 2015; Poulsen et al. 2018; but not: Edwards et al. 2006). However, to determine whether reductions in the capacity for thermal acclimation are directly caused by endocrine disruption of the TH system, future studies need to dive deeper into the pathways through which nitrate might possibly interfere with TH synthesis or signaling such as quantifying TH receptor activity in response to nitrate exposure or assessing how nitrate affects the iodine uptake by the thyroid gland. In addition to disruption of TH signaling, nitrate has been shown to impair physiological adjustments that act to increase oxygen delivery in response to higher temperatures via the remodeling of the cardio-respiratory system at all levels of the oxygen transport cascade (Gomez Isaza et al. 2020, 2021), thereby potentially reducing acclimation capacity in metabolic rate and heat tolerance.

## 5. Conclusion

Understanding how natural populations respond to global change requires consideration of exposure to multiple stressors, such as elevated temperature and toxicants, which may have adverse effects greater than those of either individual stressor (present study; Noyes et al. 2009; Rohr et al. 2011), challenging predictive power. Specifically, understanding how pollutants alter the capacity for thermal plasticity is imperative for ecological risk assessment in the context of global climate change, since acclimation capacity (Seebacher et al. 2015) as well as the capacity for temperature-induced developmental plasticity (Sinai et al. 2022) are likely to play a key role in determining species’ resilience to climate change. Given that many climate projection models predict future increases in global temperatures (IPCC 2023), and pollutants are now permanent components of the natural environment, this interaction will be of increasing relevance. If effects of a pollutant will be more pronounced under future climatic conditions, more stringent environmental quality standards will be needed for this chemical (Kimberly & Salice 2013). So far, ecological risk assessments have generally focused on single stressor exposure scenarios but data from single stressor studies rarely yield sufficient predictive power to comprehend the effects of combined stressors (Landis et al. 2013; IPCC 2023). Therefore, we highlight the need for more studies assessing the interactive effects of pollution and stressors related to climate change in order to advance predictions of population persistence under global change.

## 6. Data availability

A raw data table in xlsx format, including all original measurements, will be deposited in Figshare under DOI:XXX after acceptance.

## 8. Acknowledgements

We thank Miguel Vences, Robin Schmidt, Sven Gippner, Fabian Bartels, Cecile Zummach, and Janina Rudolph for field assistance.

## 9. Author contributions

**NS:** Data curation (supporting); Methodology (equal); Investigation (equal); Formal analysis (equal); Writing – original draft (lead). **PCE**: Data curation (supporting); Methodology (equal); Investigation (equal); Writing – review and editing (equal). **NK**: Writing – original draft (supporting); Writing – review and editing (equal). **BO**: Data curation (supporting); Investigation (equal); Formal analysis (supporting); Writing – original draft (supporting); Writing – review and editing (supporting). **KR**: Conceptualization (lead); Supervision (lead); Methodology (lead); Investigation (lead); Data curation (lead); Formal analysis (equal); Writing – original draft (lead); Funding acquisition (lead); Project administration (lead).

## 11. Statement of Ethics

The authors have no ethical conflicts to disclose. The experiments were conducted under permission from the *Niedersächsisches Landesamt für Verbraucherschutz und Lebensmittelsicherheit*, Germany (Gz. 33.19-42502-04-20/3590). Fieldwork in Lower Saxony was carried out with permits of Stadt Braunschweig (Stadt Braunschweig - Fachbereich Umwelt und Naturschutz, Richard-Wagner-Straße 1, 38106 Braunschweig; Gz. 68.11-11.8-3.3).

## 13. Supplementary material

**Table S1.** Natal pond and experimental water parameters (mean ± SD). Natal pond water parameters were determined at the time of clutch collection. Quality of tap water used in experiments was monitored before each water change twice per week during the experiment. Measurements included nitrate (NO_3_ ), nitrite (NO ^-^), ammonium (NH ^+^), pH, phosphate (PO ^3+^), copper (Cu^2+^), iron (Fe), and lead (Pb) in mg x L^-1^.

| Type | $\text{NO}_3^-$ | $\text{NO}_2^-$ | $\text{NH}_4^+$ | $\text{PO}_4^{3+}$ | $\text{Cu}^{2+}$ | Fe | Pb | pH | N |
| --- | --- | --- | --- | --- | --- | --- | --- | --- | --- |
| Natal pond water | < 6 | 0.04 ± 0.01 | 0 | 0.19 ± 0.03 | <0.02 | 0.15 ± 0.02 | 0 | 6.8 ± 0.1 | 5 |
| Tap water used in experiments | < 6 | 0 | 0 | 0.11 ± 0.01 | <0.02 | <0.1 | 0 | 7.0 ± 0.2 | 12 |

**Table S2.** Mean survival (±SD) across a range of acclimation temperatures and nitrate concentrations of the European common frog (*Rana temporaria*) from hatching until the onset of metamorphosis (Gosner stage 42; Gosner 1960).

| Nitrate concentration (mg/L) | Acclimation temperature ($^{\circ}\text{C}$) | | | |
| --- | --- | --- | --- | --- |
|  | 18 | 22 | 26 | 28 |
| 0 | 100 (±0) | 94.44 (±4.81) | 88.88 (±4.81) | 86.11 (±4.81) |
| 50 | 91.66 (±0) | 94.44 (±4.81) | 86.11 (±4.81) | 83.33 (±0) |
| 100 | 88.88 (±4.81) | 88.88 (±4.81) | 83.33 (±8.33) | 83.33 (±0) |

## 7. References

Agudelo-Cantero, G. A., & Navas, C. A. (2019). Interactive effects of experimental heating rates, ontogeny and body mass on the upper thermal limits of anuran larvae. Journal of Thermal Biology, 82, 43–51.

Albecker, M. A., Strobel, S. M., & Womack, M. C. (2023). Developmental Plasticity in Anurans: Meta-analysis Reveals Effects of Larval Environments on Size at Metamorphosis And Timing of Metamorphosis. Integrative and Comparative Biology, icad059.

Angilletta Jr, M. J., Niewiarowski, P. H., & Navas, C. A. (2002). The evolution of thermal physiology in ectotherms. Journal of Thermal Biology, 27(4), 249–268.

Angilletta, M. J. (2009). Thermal adaptation: a theoretical and empirical synthesis.

Baier, F., Jedinger, M., Gruber, E., & Zaller, J. G. (2016). Temperature-dependence of glyphosate-based herbicide’s effects on egg and tadpole growth of Common Toads. Frontiers in Environmental Science, 4, 51.

Baker, N. J., Bancroft, B. A., & Garcia, T. S. (2013). A meta-analysis of the effects of pesticides and fertilizers on survival and growth of amphibians. Science of the total environment, 449, 150–156.

Bates, D. M. (2010). lme4: Mixed-effects modeling with R.

Blaustein, A. R., & Kiesecker, J. M. (2002). Complexity in conservation: lessons from the global decline of amphibian populations. Ecology Letters, 5(4), 597–608.Blaustein et al. 2011

Boone, M. D., & Bridges, C. M. (1999). The effect of temperature on the potency of carbaryl for survival of tadpoles of the green frog (*Rana clamitans*). Environmental Toxicology and Chemistry: An International Journal, 18(7), 1482–1484.

Brooks, P. R., & Crowe, T. P. (2019). Combined effects of multiple stressors: New insights into the influence of timing and sequence. Frontiers in Ecology and Evolution, 7, 387.

Brühl, C. A., Schmidt, T., Pieper, S., & Alscher, A. (2013). Terrestrial pesticide exposure of amphibians: an underestimated cause of global decline?. Scientific reports, 3(1), 1135.

Burraco, P., & Gomez-Mestre, I. (2016). Physiological stress responses in amphibian larvae to multiple stressors reveal marked anthropogenic effects even below lethal levels. Physiological and Biochemical Zoology, 89(6), 462–472.

Claussen, D. (1977). Thermal acclimation in ambystomatid salamanders. Comparative Biochemistry and Physiology--Part A: Physiology, 58(4), 333–340.

Cowles, R. B., & Bogert, C. M. (1944). A preliminary study of the thermal requirements of desert reptiles. Bulletin of the AMNH; v. 83, article 5.

Dalvi, R. S., Pal, A. K., Tiwari, L. R., Das, T., & Baruah, K. (2009). Thermal tolerance and oxygen consumption rates of the catfish *Horabagrus brachysoma* (Günther) acclimated to different temperatures. Aquaculture, 295(1-2), 116–119.

Denver, R. J. (2021). Stress hormones mediate developmental plasticity in vertebrates with complex life cycles. Neurobiology of Stress, 14, 100301.

deWijer, P., Watt, P. J., & Oldham, R. S. (2003). Amphibian decline and aquatic pollution: effects of nitrogenous fertiliser on survival and development of larvae of the frog *Rana temporaria*. Applied Herpetology, 1, 3–12.

Díaz, S., Settele, J., Brondízio, E. S., Ngo, H. T., Agard, J., Arneth, A., … & Zayas, C. N. (2019). Pervasive human-driven decline of life on Earth points to the need for transformative change. Science, 366(6471), eaax3100.

Edwards, T. M., McCoy, K. A., Barbeau, T., McCoy, M. W., Thro, J. M., & Guillette Jr, L. J. (2006). Environmental context determines nitrate toxicity in Southern toad (*Bufo terrestris*) tadpoles. Aquatic Toxicology, 78(1), 50–58.

Eterovick, P. C., Schmidt, R., Sabino-Pinto, J., Yang, C., Künzel, S., & Ruthsatz, K. Microbiome at the Interface between Environmental Stress and Animal Health: An Example from the Most Threatened Vertebrate Group. Available at SSRN 4680902.

Garriga, N., Montori, A., & Llorente, G. A. (2017). Impact of ammonium nitrate and sodium nitrate on tadpoles of *Alytes obstetricans*. Ecotoxicology, 26, 667–674.

Goessens, T., De Baere, S., Deknock, A., De Troyer, N., Van Leeuwenberg, R., Martel, A., … & Croubels, S. (2022). Agricultural contaminants in amphibian breeding ponds: Occurrence, risk and correlation with agricultural land use. Science of the Total Environment, 806, 150661.

Gomez Isaza, D. F., Cramp, R. L., & Franklin, C. E. (2020). Living in polluted waters: A meta-analysis of the effects of nitrate and interactions with other environmental stressors on freshwater taxa. Environmental Pollution, 261, 114091.

Gomez Isaza, D. F., Cramp, R. L., & Franklin, C. E. (2021). Exposure to nitrate increases susceptibility to hypoxia in fish. Physiological and Biochemical Zoology, 94(2), 124–142.

Gomez-Mestre, I., Kulkarni, S., & Buchholz, D. R. (2013). Mechanisms and consequences of developmental acceleration in tadpoles responding to pond drying. PloS one, 8(12), e84266.

Gosner, K. L. (1960). A simplified table for staging anuran embryos and larvae with notes on identification. Herpetologica, 16(3), 183–190.

Gunderson, A. R., Armstrong, E. J., & Stillman, J. H. (2016). Multiple stressors in a changing world: the need for an improved perspective on physiological responses to the dynamic marine environment. Annual review of marine science, 8, 357–378.

Hastings, D., & Burggren, W. (1995). Developmental changes in oxygen consumption regulation in larvae of the South African clawed frog *Xenopus laevis*. Journal of experimental biology, 198(12), 2465–2475.

Hayes, T. B., Falso, P., Gallipeau, S., & Stice, M. (2010). The cause of global amphibian declines: a developmental endocrinologist’s perspective. Journal of Experimental Biology, 213(6), 921–933.

Hochachka, P. W., & Somero, G. N. (1973). Strategies of biochemical adaptation.

Holmstrup, M., Bindesbøl, A. M., Oostingh, G. J., Duschl, A., Scheil, V., Köhler, H. R., … & Spurgeon, D. J. (2010). Interactions between effects of environmental chemicals and natural stressors: a review. Science of the Total Environment, 408(18), 3746–3762.

Holzman, N., & McManus, J. J. (1973). Effects of acclimation on metabolic rate and thermal tolerance in the carpenter frog, Rana vergatipes. Comparative Biochemistry and Physiology Part A: Physiology, 45(3), 833–842.

Hooper, D. U., Adair, E. C., Cardinale, B. J., Byrnes, J. E., Hungate, B. A., Matulich, K. L., … & O’Connor, M. I. (2012). A global synthesis reveals biodiversity loss as a major driver of ecosystem change. Nature, 486(7401), 105–108.

Hourdry, J., L’Hermite, A., & Ferrand, R. (1996). Changes in the digestive tract and feeding behavior of anuran amphibians during metamorphosis. Physiological Zoology, 69(2), 219–251.

Hutchison, V. H. (1961). Critical thermal maxima in salamanders. Physiological zoology, 34(2), 92–125.

IPCC 2023: Lee, H., Calvin, K., Dasgupta, D., Krinner, G., Mukherji, A., Thorne, P., … & Park, Y. (2023). IPCC, 2023: Climate Change 2023: Synthesis Report, Summary for Policymakers. Contribution of Working Groups I, II and III to the Sixth Assessment Report of the Intergovernmental Panel on Climate Change [Core Writing Team, H. Lee and J. Romero (eds.)]. IPCC, Geneva, Switzerland.

Jaureguiberry, P., Titeux, N., Wiemers, M., Bowler, D. E., Coscieme, L., Golden, A. S., … & Purvis, A. (2022). The direct drivers of recent global anthropogenic biodiversity loss. Science advances, 8(45), eabm9982.

Johansson, M., Räsänen, K., & Merilä, J. (2001). Comparison of nitrate tolerance between different populations of the common frog, *Rana temporaria*. Aquatic Toxicology, 54(1-2), 1–14.

Kimberly, D. A., & Salice, C. J. (2013). Interactive effects of contaminants and climate-related stressors: High temperature increases sensitivity to cadmium. Environmental toxicology and chemistry, 32(6), 1337–1343.

Kingsolver, J. G., & Huey, R. B. (1998). Evolutionary analyses of morphological and physiological plasticity in thermally variable environments. American Zoologist, 38(3), 545–560.

Kirschman, L. J., McCue, M. D., Boyles, J. G., & Warne, R. W. (2017). Exogenous stress hormones alter energetic and nutrient costs of development and metamorphosis. Journal of Experimental Biology, 220(18), 3391–3397.

Kumar, R., Sharma, P., & Bandyopadhyay, S. (2021). Evidence of microplastics in wetlands: Extraction and quantification in Freshwater and coastal ecosystems. Journal of Water Process Engineering, 40, 101966.

Landis, W. G., Durda, J. L., Brooks, M. L., Chapman, P. M., Menzie, C. A., Stahl Jr, R. G., & Stauber, J. L. (2013). Ecological risk assessment in the context of global climate change. Environmental toxicology and chemistry, 32(1), 79–92.

Lannig, G., Cherkasov, A. S., & Sokolova, I. M. (2006). Temperature-dependent effects of cadmium on mitochondrial and whole-organism bioenergetics of oysters (*Crassostrea virginica*). Marine Environmental Research, 62, S79–S82.

Larsen, E. H. (2021). Dual skin functions in amphibian osmoregulation. Comparative Biochemistry and Physiology Part A: Molecular & Integrative Physiology, 253, 110869.

Laurila, A., & Kujasalo, J. (1999). Habitat duration, predation risk and phenotypic plasticity in common frog (Rana temporaria) tadpoles. Journal of Animal Ecology, 68(6), 1123–1132.

Leiva, F. P., Calosi, P., & Verberk, W. C. (2019). Scaling of thermal tolerance with body mass and genome size in ectotherms: a comparison between water-and air-breathers. Philosophical Transactions of the Royal Society B, 374(1778), 20190035.

Lind, M. I., & Johansson, F. (2007). The degree of adaptive phenotypic plasticity is correlated with the spatial environmental heterogeneity experienced by island populations of *Rana temporaria*. Journal of evolutionary biology, 20(4), 1288–1297.

Little, A. G. (2021). Thyroid hormone regulation of thermal acclimation in ectotherms: physiological mechanisms and ecoevolutionary implications. Molecular and Cellular Endocrinology, 530, 111285.

Little, A. G., & Seebacher, F. (2014). Thyroid hormone regulates cardiac performance during cold acclimation in zebrafish (*Danio rerio*). Journal of Experimental Biology, 217(5), 718–725.

Lo, L. S. H., Xu, Z., Lee, S. S., Lau, W. K., Qiu, J. W., Liu, H., … & Cheng, J. (2022). How elevated nitrogen load affects bacterial community structure and nitrogen cycling services in coastal water. Frontiers in Microbiology, 13, 1062029.

Luedtke, J. A., Chanson, J., Neam, K., Hobin, L., Maciel, A. O., Catenazzi, A., … & Stuart, S. N. (2023). Ongoing declines for the world’s amphibians in the face of emerging threats. Nature, 622(7982), 308–314.

Lutterschmidt, W. I., & Hutchison, V. H. (1997). The critical thermal maximum: data to support the onset of spasms as the definitive end point. Canadian Journal of Zoology, 75(10), 1553–1560.

Mann, R. M., Hyne, R. V., Choung, C. B., & Wilson, S. P. (2009). Amphibians and agricultural chemicals: review of the risks in a complex environment. Environmental pollution, 157(11), 2903–2927.

Merilä, J., Laurila, A., & Lindgren, B. (2004). Variation in the degree and costs of adaptive phenotypic plasticity among *Rana temporaria* populations. Journal of Evolutionary Biology, 17(5), 1132–1140.

Merilä, J., Laurila, A., Pahkala, M., Räsänen, K., & Timenes Laugen, A. (2000). Adaptive phenotypic plasticity in timing of metamorphosis in the common frog *Rana temporaria*. Ecoscience, 7(1), 18–24.

Moe, S. J., De Schamphelaere, K., Clements, W. H., Sorensen, M. T., Van den Brink, P. J., & Liess, M. (2013). Combined and interactive effects of global climate change and toxicants on populations and communities. Environmental toxicology and chemistry, 32(1), 49–61.

Morley, S. A., Peck, L. S., Sunday, J. M., Heiser, S., & Bates, A. E. (2019). Physiological acclimation and persistence of ectothermic species under extreme heat events. Global Ecology and Biogeography, 28(7), 1018–1037.

Mullur, R., Liu, Y. Y., & Brent, G. A. (2014). Thyroid hormone regulation of metabolism. Physiological Reviews, 94, 355–382

Mushtaq, N., Singh, D. V., Bhat, R. A., Dervash, M. A., & Hameed, O. B. (2020). Freshwater contamination: sources and hazards to aquatic biota. Fresh water pollution dynamics and remediation, 27–50.

Navas, C. A., Gomes, F. R., & Carvalho, J. E. (2008). Thermal relationships and exercise physiology in anuran amphibians: integration and evolutionary implications. Comparative Biochemistry and Physiology Part A: Molecular & Integrative Physiology, 151(3), 344–362.

Newman, R. A. (1992). Adaptive plasticity in amphibian metamorphosis. BioScience, 42(9), 671–678.

Noyes, P. D., & Lema, S. C. (2015). Forecasting the impacts of chemical pollution and climate change interactions on the health of wildlife. Current Zoology, 61(4), 669–689.

Noyes, P. D., McElwee, M. K., Miller, H. D., Clark, B. W., Van Tiem, L. A., Walcott, K. C., … & Levin, E. D. (2009). The toxicology of climate change: environmental contaminants in a warming world. Environment international, 35(6), 971–986.

Orlofske, S. A., & Hopkins, W. A. (2009). Energetics of metamorphic climax in the pickerel frog (Lithobates palustris). Comparative Biochemistry and Physiology Part A: Molecular & Integrative Physiology, 154(2), 191–196.

Ortiz-Santaliestra, M. E., & Sparling, D. W. (2007). Alteration of larval development and metamorphosis by nitrate and perchlorate in southern leopard frogs (*Rana sphenocephala*). Archives of environmental contamination and toxicology, 53, 639–646.

Osterauer, R., & Köhler, H. R. (2008). Temperature-dependent effects of the pesticides thiacloprid and diazinon on the embryonic development of zebrafish (*Danio rerio*). Aquatic Toxicology, 86(4), 485–494.

Pirotta, E., Thomas, L., Costa, D. P., Hall, A. J., Harris, C. M., Harwood, J., … & Tyack, P. L. (2022). Understanding the combined effects of multiple stressors: A new perspective on a longstanding challenge. Science of the Total Environment, 821, 153322.

Poulsen, R., Cedergreen, N., Hayes, T., & Hansen, M. (2018). Nitrate: an environmental endocrine disruptor? A review of evidence and research needs. Environmental science & technology, 52(7), 3869–3887.

Pulsford, S. A., Barton, P. S., Driscoll, D. A., & Lindenmayer, D. B. (2019). Interactive effects of land use, grazing and environment on frogs in an agricultural landscape. Agriculture, Ecosystems & Environment, 281, 25–34.

R Core Team. (2021). *R: A Language and Environment for Statistical Computing*. Vienna, Austria. Retrieved from https://www.R-project.org/

Rohr, J. R., & Palmer, B. D. (2005). Aquatic herbicide exposure increases salamander desiccation risk eight months later in a terrestrial environment. Environmental Toxicology and Chemistry: An International Journal, 24(5), 1253–1258.

Rohr, J. R., & Palmer, B. D. (2013). Climate change, multiple stressors, and the decline of ectotherms. Conservation Biology, 27(4), 741–751.

Rohr, J. R., Kerby, J. L., & Sih, A. (2006). Community ecology as a framework for predicting contaminant effects. Trends in Ecology & Evolution, 21(11), 606–613.

Rohr, J. R., Sesterhenn, T. M., & Stieha, C. (2011). Will climate change reduce the effects of a pesticide on amphibians?: partitioning effects on exposure and susceptibility to contaminants. Global Change Biology, 17(2), 657–666.

Rouse, J. D., Bishop, C. A., & Struger, J. (1999). Nitrogen pollution: an assessment of its threat to amphibian survival. Environmental health perspectives, 107(10), 799–803.

Ruthsatz, K., & Glosv, J. (2024). Effects of Pollutants on the Endocrine System of Tadpoles. Toxicology of Amphibian Tadpoles, 78–110.

Ruthsatz, K., Bartels, F., Stützer, D., & Eterovick, P. C. (2022a). Timing of parental breeding shapes sensitivity to nitrate pollution in the common frog *Rana temporaria*. Journal of Thermal Biology, 108, 103296.

Ruthsatz, K., Dausmann, K. H., Drees, C., Becker, L. I., Hartmann, L., Reese, J., … & Glos, J. (2020a). Altered thyroid hormone levels affect the capacity for temperature-induced developmental plasticity in larvae of *Rana temporaria* and *Xenopus laevis*. Journal of Thermal Biology, 90, 102599.

Ruthsatz, K., Dausmann, K. H., Paesler, K., Babos, P., Sabatino, N. M., Peck, M. A., & Glos, J. (2020b). Shifts in sensitivity of amphibian metamorphosis to endocrine disruption: the common frog (*Rana temporaria*) as a case study. Conservation Physiology, 8(1), coaa100.

Ruthsatz, K., Dausmann, K. H., Peck, M. A., & Glos, J. (2022b). Thermal tolerance and acclimation capacity in the European common frog (*Rana temporaria*) change throughout ontogeny. Journal of Experimental Zoology Part A: Ecological and Integrative Physiology, 337(5), 477–490.

Ruthsatz, K., Dausmann, K. H., Peck, M. A., Drees, C., Sabatino, N. M., Becker, L. I., … & Glos, J. (2018a). Thyroid hormone levels and temperature during development alter thermal tolerance and energetics of *Xenopus laevis* larvae. Conservation Physiology, 6(1), coy059.

Ruthsatz, K., Dausmann, K. H., Reinhardt, S., Robinson, T., Sabatino, N. M., Peck, M. A., & Glos, J. (2019). Endocrine disruption alters developmental energy allocation and performance in *Rana temporaria*. Integrative and Comparative Biology, 59(1), 70–88.

Ruthsatz, K., Dausmann, K. H., Reinhardt, S., Robinson, T., Sabatino, N. M., Peck, M. A., & Glos, J. (2020c). Post-metamorphic carry-over effects of altered thyroid hormone level and developmental temperature: physiological plasticity and body condition at two life stages in Rana temporaria. Journal of Comparative Physiology B, 190, 297–315.

Ruthsatz, K., Peck, M. A., Dausmann, K. H., Sabatino, N. M., & Glos, J. (2018b). Patterns of temperature induced developmental plasticity in anuran larvae. Journal of Thermal Biology, 74, 123–132.

Ruthsatz, K., Schwarz, A., Gomez-Mestre, I., Meyer, R., Domscheit, M., Bartels, F., … & Engelkes, K. (2023). Life in plastic, it’s not fantastic: Sublethal effects of polyethylene microplastics ingestion throughout amphibian metamorphosis. Science of The Total Environment, 885, 163779.

Rzymski, P., Drewek, A., & Klimaszyk, P. (2017). Pharmaceutical pollution of aquatic environment: An emerging and enormous challenge. Limnological Review, 17(2), 97–107.

Sala, O. E., Stuart Chapin, F. I. I. I., Armesto, J. J., Berlow, E., Bloomfield, J., Dirzo, R., … & Wall, D. H. (2000). Global biodiversity scenarios for the year 2100. science, 287(5459), 1770–1774.

Seebacher, F., White, C. R., & Franklin, C. E. (2015). Physiological plasticity increases resilience of ectothermic animals to climate change. Nature Climate Change, 5(1), 61–66.

Shi, Y. B. (2000). *Amphibian metamorphosis: from morphology to molecular biology* (No. 19638). Wiley-Liss.

Sigmund, G., Ågerstrand, M., Antonelli, A., Backhaus, T., Brodin, T., Diamond, M. L., … & Groh, K. J. (2023). Addressing chemical pollution in biodiversity research. Global Change Biology, 29(12), 3240–3255.

Sih, A., Bell, A. M., & Kerby, J. L. (2004). Two stressors are far deadlier than one. Trends in ecology & evolution, 19(6), 274–276.

Sinai, N., Glos, J., Mohan, A. V., Lyra, M. L., Riepe, M., Thöle, E., … & Ruthsatz, K. (2022). Developmental plasticity in amphibian larvae across the world: Investigating the roles of temperature and latitude. Journal of Thermal Biology, 106, 103233.

Sundermann, G., Wägner, N., Cullmann, A., von Hirschhausen, C. R., & Kemfert, C. (2020). Nitrate pollution of groundwater long exceeding trigger value: Fertilization practices require more transparency and oversight. DIW Weekly Report, 10(8/9), 61–72.

Tiso, M., & Schechter, A. N. (2015). Nitrate reduction to nitrite, nitric oxide and ammonia by gut bacteria under physiological conditions. PloS one, 10(3), e0119712.

Trudeau, V. L., Thomson, P., Zhang, W. S., Reynaud, S., Navarro-Martin, L., & Langlois, V. S. (2020). Agrochemicals disrupt multiple endocrine axes in amphibians. Molecular and Cellular Endocrinology, 513, 110861.

Turriago, J. L., Parra, C. A., & Bernal, M. H. (2015). Upper thermal tolerance in anuran embryos and tadpoles at constant and variable peak temperatures. Canadian Journal of Zoology, 93(4), 267–272.

Van Meter, R. J., Adelizzi, R., Glinski, D. A., & Henderson, W. M. (2019). Agrochemical mixtures and amphibians: the combined effects of pesticides and fertilizer on stress, acetylcholinesterase activity, and bioaccumulation in a terrestrial environment. Environmental toxicology and chemistry, 38(5), 1052–1061.

Vences, M., Sarasola-Puente, V., Sanchez, E., Amat, F., & Hauswaldt, J. S. (2017). Diversity and distribution of deep mitochondrial lineages of the common frog, *Rana temporaria*, in northern Spain. Salamandra, 53(1).

von May, R., Catenazzi, A., Santa-Cruz, R., Gutierrez, A. S., Moritz, C., & Rabosky, D. L. (2019). Thermal physiological traits in tropical lowland amphibians: Vulnerability to climate warming and cooling. PLoS One, 14(8), e0219759.

Wake, D. B., & Vredenburg, V. T. (2008). Are we in the midst of the sixth mass extinction? A view from the world of amphibians. Proceedings of the National Academy of Sciences, 105(supplement_1), 11466–11473.

Walls, S. C., & Gabor, C. R. (2019). Integrating behavior and physiology into strategies for amphibian conservation. Frontiers in Ecology and Evolution, 7, 234.

Wang, M., Chai, L., Zhao, H., Wu, M., & Wang, H. (2015). Effects of nitrate on metamorphosis, thyroid and iodothyronine deiodinases expression in *Bufo gargarizans* larvae. Chemosphere, 139, 402–409.

Wu, C. S., & Kam, Y. C. (2005). Thermal tolerance and thermoregulation by Taiwanese rhacophorid tadpoles (*Buergeria japonica*) living in geothermal hot springs and streams. Herpetologica, 61(1), 35–46.

Wu, N. C., & Seebacher, F. (2021). Bisphenols alter thermal responses and performance in zebrafish (*Danio rerio*). Conservation Physiology, 9(1).

Xie, L., Zhang, Y., Qu, Y., Chai, L., Li, X., & Wang, H. (2019). Effects of nitrate on development and thyroid hormone signaling pathway during *Bufo gargarizans* embryogenesis. Chemosphere, 235, 227–238.

